# Female genetic contributions to sperm competition in *Drosophila melanogaster*

**DOI:** 10.1101/500546

**Authors:** Dawn S. Chen, Sofie Y.N. Delbare, Simone L. White, Jessica Sitnik, Martik Chatterjee, Elizabeth DoBell, Orli Weiss, Andrew G. Clark, Mariana F. Wolfner

## Abstract

In many species, sperm can remain viable in the reproductive tract of a female well beyond the typical interval to remating. This creates an opportunity for sperm from different males to compete for oocyte fertilization inside the female’s reproductive tract. In *Drosophila melanogaster*, sperm morphology and seminal fluid content affect male success in sperm competition. On the other hand, although genome-wide association studies (GWAS) have demonstrated that female genotype plays a role in sperm competition outcome as well, the biochemical, sensory and physiological processes by which females detect and selectively use sperm from different males remain elusive.

Here, we functionally tested 27 candidate genes implicated via a GWAS for their contribution to the female’s role in sperm competition, measured as changes in the relative success of the first male to mate (P1). Of these 27 candidates, we identified eight genes that affect P1 when knocked down in females, and also showed that six of them do so when knocked down in the female nervous system. Two genes in particular, *Rim* and *caup*, lowered P1 when knocked down in sensory *pickpocket* (*ppk*)^+^ neurons and octopaminergic *Tdc2*^+^ neurons, respectively. These results establish a functional role for the female’s nervous system in the process of sperm competition and expand our understanding of the genetic, neuronal and mechanistic basis of female responses to multiple matings. We propose that through their nervous system, females actively assess male compatibility based on courtship or ejaculates and modulate sperm competition outcome accordingly.

## Introduction

Natural and sexual selection increase the frequencies of alleles that boost an organism’s reproductive success. Sexual selection acts on pre-copulatory traits, such as male courtship behavior and female mate choice, as well as on post-copulatory processes. Across vertebrates and invertebrates, it can be beneficial for females to obtain multiple mates (Jennions and Petrie 2000). If multiple mating occurs at a high enough frequency, and/or if sperm is stored long term, ejaculates from rival males will compete for oocyte fertilization (Parker 1970). This type of male-male post-copulatory sexual selection mediates the evolution of adaptations in males to cope with the opportunities for sperm competition. One form of adaptation is to lower the chances of female re-mating with other males, since the last male to mate often sires most of a female’s progeny (Chen et al. 1988, Orr and Rutowski 1991, Baer et al. 2001, Birkhead and Pizzari 2002, Chapman et al. 2003, Liu and Kubli 2003, Wedell 2005, Wigby and Chapman 2005, Laturney and Billeter 2016, Sutter and Lindholm 2016).

If a female does re-mate, characteristics of sperm and seminal fluid proteins influence a male’s ability to compete with ejaculates from other males. *Drosophila melanogaster* has proven to be an especially informative model to study these male × male genotypic interactions. Generally, longer and slower sperm are better at withstanding displacement in *D. melanogaster* (Lüpold et al. 2012). Genome-wide association studies (GWAS) further uncovered the genetic basis of male competitive ability. Besides genes encoding sperm components (Yeh et al. 2012), genes encoding seminal fluid proteins were discovered to play a role in sperm competition (Clark et al. 1995, Fiumera et al. 2005, Fiumera et al. 2007, Greenspan and Clark 2011). These proteins have a variety of functions, such as inducing female refractoriness to re-mating and stimulating egg laying (e.g. Sex peptide; Chapman et al. 2003) and promoting sperm storage (e.g. Acp36DE (Neubaum and Wolfner 1999), Acp29AB (Wong et al. 2008), and Acp62F (Mueller et al. 2008)). Interestingly, many seminal fluid proteins evolve rapidly (reviewed in Swanson and Vacquier 2002), and some were found to be harmful to females (Civetta and Clark 2000, Lung et al. 2001, Wigby and Chapman 2005, Mueller et al. 2007), suggesting that their evolution is mediated by sexual conflict: what makes a male a better competitor might actually be disadvantageous to females.

Although most studies of sperm competition focused on the roles of the male, a number of studies have argued that females are not “passive vessels” in this process. Cryptic female choice, whereby a female selectively uses sperm from ejaculates she received from multiple males, has been proposed as a powerful mechanism for female contributions to sperm competition (Eberhard 1996). A classic example of such female contribution has been observed in junglefowl, in which females were seen to eject sperm from subdominant males after forced copulation (Pizzari and Birkhead 2000). Studies in *D. melanogaster*, with standard male genotypes and varying female genotypes also illustrate that male success depends not only on his genotype and the genotype of his competitor, but also on the genotype of the female (Clark et al. 1999, Clark et al. 2000, Lawniczak and Begun 2005, Chow et al. 2010, Giardina et al. 2011, Lüpold et al. 2013, Zhang et al. 2013, Reinhart et al. 2015). These three-way interactions have been suggested to be important for maintaining polymorphisms in populations (Clark et al. 2000, Clark 2002). However, despite the observation that female genotype plays a role, it has been difficult to disentangle female control from female × male interactions, and to demonstrate the genetics of this control. Recent studies in *Drosophila* have begun to provide a way to dissect the female’s role in sperm competition, and to determine the genes and mechanisms that contribute to differences in sperm competition outcome. First, *D. melanogaster* males carrying sperm protamines labeled with GFP or RFP enabled direct observation of competing sperm inside the female reproductive tract (Manier et al. 2010) and measurements of heritable variation across female genotypes in sperm ejection, storage and displacement (Lüpold et al. 2013). Second, initial studies have been done of the female’s genetic makeup underlying variation in her contribution to sperm competition. Chow et al. (2013) identified SNPs whose presence in the female was associated with sperm competition outcome, by performing sperm competition assays using two standard tester males and females from 39 DGRP lines, a panel of wild-derived inbred lines whose genome sequences are available (Mackay et al. 2012). They found variation in the relative number of first male offspring (P1) across DGRP females, and a GWAS revealed correlations between P1 and SNPs in or close to 33 genes (Chow et al. 2013). However, roles for these genes in sperm competition were not known. Fifteen of the 33 candidate genes identified by SNPs by Chow et al. (2013) have expression biased to the nervous system, or have known neural functions, encoding proteins such as ion channels, transcription factors involved in proneural development, or proteins with roles in vesicle trafficking. This led Chow et al. to suggest a role for the female nervous system in impacting the paternity share of each male. The proposal that the female nervous system might play a role in sperm competition is further supported by findings regarding sex peptide receptor (*SPR;* Chow et al. 2010) and *Neprilysin 2* (Sitnik et al. 2014), which to date are the only two genes known to affect female contributions to sperm competition. This was determined in experiments that knocked down SPR or Neprilysin 2 in females ubiquitously, but both genes are known to be expressed in the female nervous system.

Although not previously shown to impact sperm competition, the female nervous system is known to play important roles in post-mating responses (Arthur et al. 1998). Activity of a neuronal circuit involving *Dh44*^+^ neurons has been shown to influence the time after mating at which females eject sperm, a process that can influence competitive fertilization success (Lee et al. 2015). The neuromodulator octopamine, as well as octopaminergic *Tdc2*^+^ neurons, are required for sperm release from storage (Avila et al. 2012), ovulation (Rubinstein and Wolfner 2013), and refractoriness to re-mating (Häsemeyer et al. 2009, Rezával et al. 2014). Furthermore, subsets of sensory *pickpocket* (*ppk*)^+^ neurons that express *SPR* are required for increased egg laying and refractoriness post-mating (Yapici et al. 2008, Yang et al. 2009, Rezával et al. 2012).

Here, we performed functional tests to determine directly whether any of the candidate genes put forward by Chow et al. (2013) affect sperm competition and, for those that did, whether they elicit their effects through the female’s nervous system. We individually knocked down candidate genes using RNA interference (RNAi) in females, either ubiquitously or in the nervous system. Knockdown and control females were mated consecutively to two distinct tester males and we assessed the effect of knockdown on paternity ratios. Of 27 genes tested, eight genes were found to affect the ratio of offspring sired by each male. Having identified genes that are essential in females for sperm competition outcome, we then tested whether their role was in the nervous system as a whole, or, because of their roles in modulating female post-mating responses, in *Tdc2*^+^ and *ppk*^+^ neurons.

Six of these eight genes affected sperm competition outcome when knocked down in the entire female nervous system, or in *Tdc2*^+^ or *ppk*^+^ neurons. Our results provide the first proof that the female plays an active role in sperm competition, via action of particular genes in her nervous system. We also identified two subsets of neurons with involvement in this process. These results will allow detailed dissection of the mechanisms of cryptic female choice and sperm competition inside the female reproductive tract, and by extension effects of post-mating pre-zygotic sexual selection and sexual conflict.

## Materials and Methods

### Fly stocks and husbandry

The UAS/GAL4 system (Brand and Perrimon 1993) was used to individually knock down candidate genes ubiquitously, pan-neuronally or in subsets of the female nervous system. Driver lines used were: ubiquitous drivers *Tubulin*-GAL4/TM3, *Sb* and *Tubulin*-GAL80^ts^; *Tubulin*-GAL4/TM3, *Sb*, nervous system-specific drivers *nSyb*-GAL4 (Hindle et al. 2013), *ppk*-GAL4 (Matthews et al. 2007) and *Tdc2*-GAL4 (Cole et al. 2005). UAS-RNAi lines were ordered from the Vienna *Drosophila* Research Center (VDRC) for each candidate gene identified in a GWAS (Chow et al. 2013) with the following exceptions: *CG10858, RFeSP* and *sti* (no VDRC lines were available for these genes), and *CG13594* (the only available VDRC line has 94 predicted off-targets). VDRC IDs for all VDRC lines are available in table S1. Lines were used from both the *attP* and *w^1118^* backgrounds. To obtain controls with wild-type gene expression, flies from the appropriate background stock were crossed with flies from the driver lines. Males used for the sperm competition assay had the *cn bw* or *bw^D^* genotypes (see below). Males and virgin knockdown and control females were aged 3-7 days in single-sex vials before the start of each experiment.

Fly stocks were maintained at room temperature on standard yeast/glucose media on a 12 hr-light/dark cycle. When using *Tubulin*-GAL80^ts^; *Tubulin*-GAL4/TM3, *Sb*, crosses were set up at room temperature, and knockdown and control virgin females were aged at 29°C and maintained at 29°C throughout the sperm competition assay.

### Verification of knockdown efficiency

To verify knockdown efficiency, UAS-RNAi lines were crossed to *Tubulin*-GAL4/*TM3, Sb.* Five candidate genes did not yield viable *Tubulin*-GAL4>UAS-RNAi F1 progeny, suggesting that ubiquitous knockdown of the target gene was lethal and that the RNAi-mediated knockdown was efficient. For crosses that yielded viable *Tubulin*-GAL4>UAS-RNAi F1 progeny, RT-PCR was used to assess knockdown efficiency of each UAS-RNAi line, as previously described (Ravi Ram et al. 2006; Table S1; primers available upon request). Age-matched *TM3, Sb;* UAS-RNAi siblings or *Tubulin-*GAL4>*w^1118^* or *Tubulin*-GAL4>*attP* flies were collected at the same time as knockdown flies and tested as controls.

### Sperm competition experiments

Control and knockdown females were mated to *cn bw* males in single pair matings on day 0 in vial 1. Copulations were observed. Males were removed after copulation ended and mated females retained in the individual vials. In the evening of day 1, two *bw^D^* males were added to each vial and left with the female overnight. Both *bw^D^* males were removed in the morning of day 2, and each female was transferred to vial 2. Each female was transferred again every 48 hrs to vials 3, 4 and 5 (on days 4, 6 and 8, respectively). All females were discarded on day 10. Progeny from eggs laid in vials 1-5 were reared to adulthood and the paternity of F1 female progeny was scored based on eye color: female offspring of *cn bw* males had red eyes, and female offspring of *bw^D^* males had brown eyes. Male progeny were not scored because they were *w*^-^, making it impossible to use eye color to assess their paternity.

On average, each experiment consisted of 71.8 ± 25.1 control females and 65.9 ± 24.3 knockdown females who had mated at least once. Of these females, 51.9 ± 21.7 control females and 46.9 ± 21.0 knockdown females in each experiment had mated with both males (Table S2). All sample sizes are reported in mean ± standard deviation.

### Statistical analysis of remating rate, fertility and P1

Remating rate, fertility, and P1 of knockdown and control females were calculated based on the numbers of first- and second-male progeny. All statistical analyses were performed using base R (version 3.3.1; R Core Team 2016) and the packages lme4 (Bates et al. 2015), lmerTest (Kuznetsova et al. 2017) and emmeans (https://cran.r-project.org/web/packages/emmeans/index.html).

Remating rate was calculated as the proportion of doubly-mated females among all females who mated with the first male. Differences between remating rates of knockdown and control females were compared using Fisher’s exact test.

Because we only scored eye color in female offspring, we used the total number of female progeny produced by each doubly-mated female (rather than all progeny) as a proxy for fertility. Females who had mated only once (with the first or the second male, but not both males) were excluded from the analysis because we found a significant study-wide difference between the fertility of singly-mated and doubly-mated females (Fig. S1). We compared fertility of control and knockdown females by fitting linear models, or linear mixed models for experiments with multiple replicates, and comparing the estimated marginal means. To compare the temporal dynamics of fertility in vials 1-5 between control and knockdown females, we used linear mixed models with genotype and vial as fixed effects and individual females as a random effect.

Finally, P1 was calculated for each doubly-mated female as the ratio of the number of female offspring sired by the first male vs. the total number of female offspring sired by either the first or second males in vials 2-5. Vial 1 was excluded from the calculation of P1 because both matings occurred in vial 1, and with this experimental setup, we were unable to determine how many offspring were sired before the second mating. However, the presence of first- and/or second-male progeny in all vials was used to determine whether a female had mated with both males. For the statistical analysis of P1, we arcsine square-root transformed P1 values before applying linear models or linear mixed models for experiments with multiple replicates to the transformed values. Temporal dynamics of P1 between control and knockdown females were also compared using linear mixed models, with genotype and vial as fixed effects and individual females as random effect.

## Results

Chow et al. (2013) identified 33 top SNPs that associated with sperm competition outcome in females. Not all of these SNPs were located within genes. Thus, to identify genes that directly affect sperm competition, we used RNAi to individually knock down genes that were put forward as candidates by Chow et al. We tested 27 of the 33 candidate genes for roles in influencing the female’s contribution to sperm competition, which we scored as P1, the proportion of first male progeny among total progeny after the second mating. Four of the genes that were identified by Chow et al. could not be tested because no suitable UAS-RNAi lines were available from the VDRC. RNAi lines for two additional genes, *SK* and *CG33298*, gave insufficient RNAi knockdown for testing (Table S1), and thus, we were unable to assess these two genes’ role in sperm competition.

### Candidate gene knockdown caused changes in fertility and remating rate

How readily females remate after the first mating and how fertile they are can influence the risk and intensity of sperm competition. Therefore, we assessed the effect of knocking down each of the 27 genes on remating rate and fertility. Of the 27 genes, ubiquitous or tissue-specific knockdown of 6 genes reduced female remating rate (*CG10962, CG33095, Ddr, para, Rab2, spz;* Table 1, Fig. S2), and *Tdc2*^+^ and *ppk*^+^ neuron-specific knockdown of *hid* led to an increase in remating rate (Table 1, Fig. S2). Since *hid* expression stimulates apoptosis (Grether et al. 1995), differences in the numbers or innervation patterns of *Tdc2*^+^ and *ppk*^+^ neurons might be responsible for this effect. Reduced remating rate observed upon ubiquitous knockdown needs to be interpreted with caution, since ubiquitous knockdowns could directly affect female receptivity to remating, or could have detrimental effects on overall female health or development, making females simply less inclined to mate.

**Table 1:** Effects of gene knockdown (KD) on female remating rate. Asterisks indicate p < 0.05 (*), p < 0.01 (**) or p < 0.001 (***).

| Gene(s) | Driver(s) | Effect of KD on remating rate | p-value |
| --- | --- | --- | --- |
| *CG10962*<br>*CG33095*<br>*Ddr*<br>*spz* | *Tubulin* | ↓ | ***<br>*<br>**<br>* |
| *Rab2* | *Tubulin; Tdc2<sup>+</sup>* | ↓ | *** |
| *hid* | *Tdc2<sup>+</sup>; ppk<sup>+</sup>* | ↑ | *** |

Female fertility was affected by many candidate gene knockdowns. Ubiquitous knockdown of 19 of the 27 genes reduced female fertility (Table 2, Fig. S2). However, as mentioned above, these results could be either direct or indirect consequences of ubiquitous gene knockdown. Consistent with the latter hypothesis, we found that nervous system-specific knockdown of only 5 genes caused a decrease in female fertility *(btsz, caup, Ddr, Rab2, Rim;* Table 2, Fig. S2). Specifically, *Tdc2*^+^ neuron-specific knockdown of *Rab2* mediated a substantial decrease in both fertility and remating rate, to the extent that very few doubly mated females were retrieved for sperm competition experiments (only 8 out of 62 females remated). Because the knockdown was tissue specific, these results strongly suggest that *Rab2* is essential for the proper functioning of *Tdc2*^+^ neurons, which are in turn known to be required for female remating and fecundity (Rezával et al. 2014). Interestingly, ubiquitous knockdown of *Zasp66* and *ppk*^+^ neuron-specific knockdown of *hid* significantly increased fertility (Table 2, Fig. S2).

**Table 2:** Effects of gene knockdown (KD) on female fertility. Asterisks indicate p < 0.05 (*), p < 0.01 (**) or p < 0.001 (***).

| Gene(s) | Driver(s) | Effect of KD on fertility | p-value |
| --- | --- | --- | --- |
| *Ddr* | *ppk<sup>+</sup>*<br>*Tdc2<sup>+</sup>*<br>*nSyb* | ↓ | **<br>*<br>*** |
| *Rim* | *Tdc2<sup>+</sup>* | ↓ | * |
| *Rab2* | *Tdc2<sup>+</sup>* | ↓ | *** |
| *caup* | *Tdc2<sup>+</sup>* | ↓ | * |
| *btsz* | *nSyb*<br>*Tubulin* | ↓ | *<br>** |
| *Zasp66* | *Tubulin* | ↑ | *** |
| *hid* | *ppk<sup>+</sup>* | ↑ | *** |
| *CG10962*;<br>*CG15800*;<br>*CG31872*;<br>*CG32532*;<br>*CG32834*;<br>*CG33095*;<br>*CG33298*;<br>*CG42796*;<br>*CG6163*; *CG9850*;<br>*Cyp313a2*;<br>*Msp300*; *Rbp6*;<br>*Shab*; *sima*; *spz*;<br>*spz5*; *uif* | *Tubulin* | ↓ | Range *-*** |

Although all 27 candidate genes were detected in a GWAS based on sperm competition outcomes, these results suggest that some of the genes play roles in modulating other female reproductive traits. In particular, they suggest that fertility and sperm competition outcome should not be regarded as isolated results, since they both depend on a female’s reproductive output. Therefore, when reporting P1 below, we also report any differences in absolute first- and second-male progeny counts between control and knockdown females.

### Seven genes influence sperm competition outcome upon ubiquitous or pan-neuronal knockdown in females

Of the 27 candidate genes of interest, three *(para, Rim* and *Rab2*) were reported to affect P1 when knocked down in ppk^+^ neurons by Chow et al. (2013). For the 24 remaining candidate genes, in an initial test we knocked down each candidate ubiquitously with *Tubulin*-GAL4. If constitutive ubiquitous knockdown was lethal, and/or if the gene of interest had a known neural function, *Tubulin*-GAL4; *Tubulin*-GAL80^ts^ or the pan-neuronal driver nSyb-GAL4 were used instead of *Tubulin*-GAL4. In cases where ubiquitous knockdown produced a significant effect on overall P1, we proceeded to knock down the gene pan-neuronally, with the exception of *CG31872* and *CG32834.* These two genes are not expressed in the nervous system, but are expressed in the female rectal pad and sperm storage organs, respectively (Leader et al. 2018). We hypothesize that the effects of knockdown on sperm competition outcome may be due to the importance of these genes’ expression in the female reproductive tract.

Ubiquitous knockdown of five genes in females caused reduction of P1 (*btsz, CG31872, CG32834, Ddr, Msp300;* Table 3, Fig. S2), attributable to fewer first male progeny and more second male progeny (*CG31872* and *CG32834*, Fig. 1B, C), fewer first and second male progeny (*btsz, Msp300*, Fig. 1A, E), or fewer first male progeny but similar numbers of second male progeny relative to control females (*Ddr*, Fig. 1D). The overall fertility of these knockdown females was also lower than that of control females for all five genes.

**Table 3:** Effects of gene knockdown (KD) on sperm competition outcome. Sperm competition outcome was measured as the relative number of offspring sired by the first male (P1). Asterisks indicate p < 0.05 (*), p < 0.01 (**) or p < 0.001 (***).

| **Gene** | **Driver(s)** | **Effect of KD on P1** | **P-value** |
| --- | --- | --- | --- |
| *CG31872* | *Tubulin* | ↓ | ** |
| *CG32834* | *Tubulin* | ↓ | *** |
| *Ddr* | *Tubulin* | ↓ | * |
| *btsz* | *Tubulin*<br>*nSyb* | ↓<br>↓ | *<br>*** |
| *Caup* | *nSyb*<br>*Tdc2<sup>+</sup>* | ↓<br>↓ | **<br>** |
| *hid* | *nSyb* | ↑ | * |
| *Msp300* | *Tubulin*<br>*nSyb* | ↓<br>↑ | ***<br>* |
| *Rim* | *ppk<sup>+</sup>* | ↓ | * |

**Fig 1.**
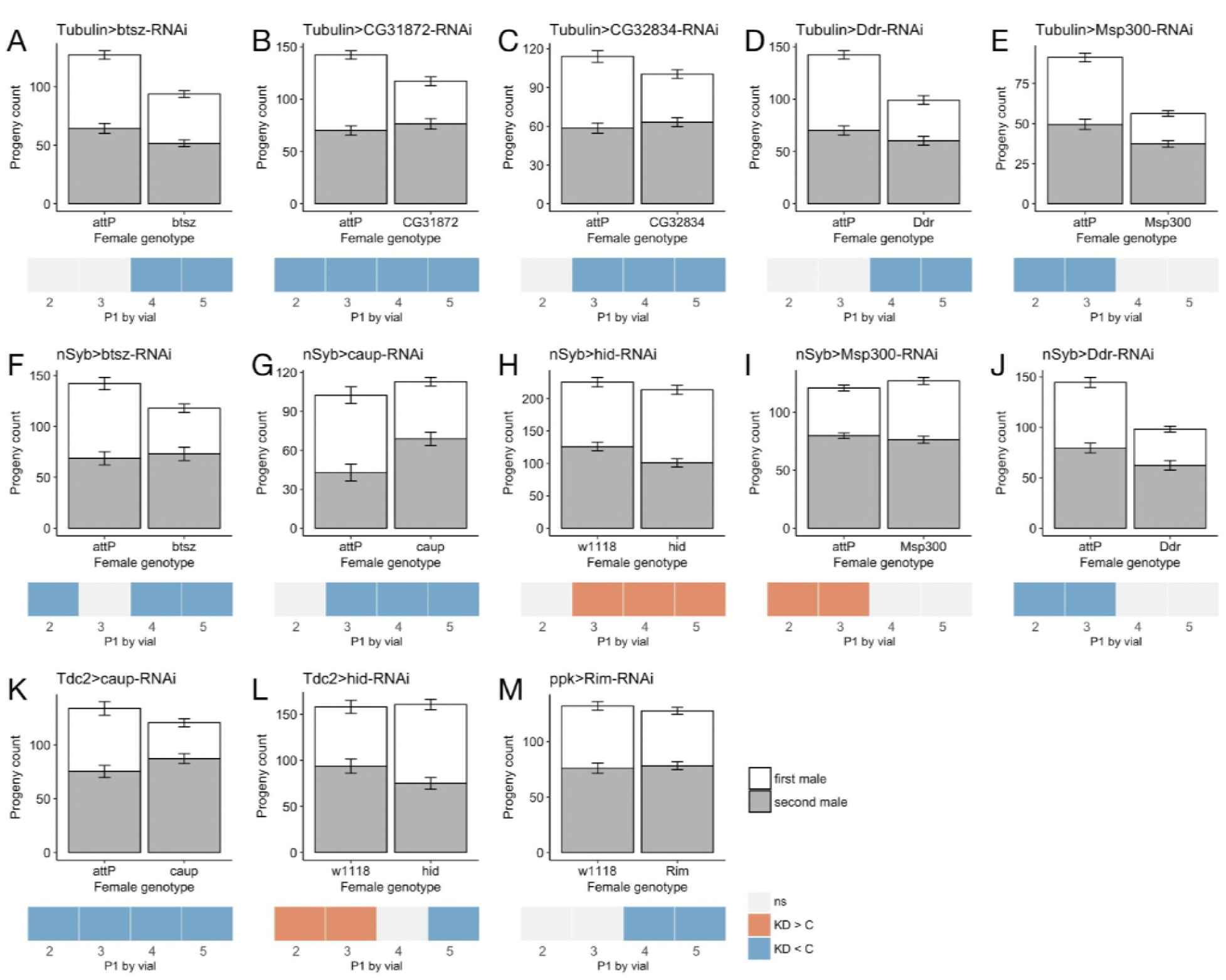
Tissue-specific knockdown of eight genes affects sperm competition, and in some cases, female fertility. Results for genes with significant effects on fertility and/or P1 when knocked down ubiquitously (A-E), pan-neuronally (F-J), in *Tdc2*^+^ neurons (K, L) or in *ppk*^+^ neurons (M) are shown. Barplots represent mean ± SEM of female progeny counts sired by the first (white) and second male (gray). Heatmaps show the temporal (per vial) differences in P1 between knockdown and control females. Colored boxes indicate significant differences, either knock-down females having higher P1 than control females (KD > C; orange) or knock-down females having lower P1 than control females (KD < C; blue).

Additionally, we found four genes whose pan-neuronal knockdown caused an increase (*hid, Msp300*) or decrease (*btsz, caup*) in P1 (Table 3, Fig. S2). Pan-neuronal knockdown of *btsz* reduced the number of first male progeny without affecting the number of second male progeny, leading to an overall reduction in fertility (Fig. 1F). Pan-neuronal knockdown of *caup* and *hid* affected the relative proportions of first- and second-male progeny without influencing overall fertility (Fig. 1G, H). Finally, *Msp300* pan-neuronal knockdown females produced more first male progeny but similar numbers of second male progeny compared to control females, but the overall fertility difference between Msp300 knockdown and control females was not significant (Fig. 1I). Intriguingly, ubiquitous knockdown of *Msp300* lowered P1, while pan-neuronal knockdown increased P1 (Table 3, Fig. S2). This suggests that ubiquitous knockdown of *Msp300* could be detrimental to females’ health, or that *Msp300* expression in different tissues has distinct effects on sperm competition. Overall, we found eight genes that had effects on sperm competition when knocked down ubiquitously or pan-neuronally in females.

When analyzing P1 on a temporal, vial by vial basis, we found that at least two vials were significantly different between control and knockdown females for each of the genes that had an effect on overall P1 (Fig. 1). *Ddr*, which affected overall P1 upon ubiquitous knockdown only, also showed significant effects on P1 in vials 2 and 3 with pan-neuronal knockdown (Fig. 1J), suggesting some neuronal function for *Ddr* as well. Finally, three genes (*sima, spz5* and *Zasp66*) did not change overall P1 when knocked down ubiquitously, but significantly affected P1 in 2 vials when ubiquitous knockdown was analyzed on a vial by vial basis (Fig. S3). This suggests that the products of these three genes might be important for sperm competition at specific times after the second mating. Alternatively, these gene products or the processes they mediate might have small or redundant roles in sperm competition.

### *Tdc2*^+^ and *ppk*^+^ neurons play roles in sperm competition

Informed by the results of the initial test, we further asked in which of the female’s neurons the products of *btsz, caup, hid, Msp300* and *Ddr* act to modulate sperm competition. In particular, we assessed the functions of these five genes in octopaminergic *Tdc2*^+^ neurons and proprioceptive *ppk*^+^ neurons, which have known roles in female responses to mating (Cole et al. 2005, Yapici et al. 2008, Häsemeyer et al. 2009, Yang et al. 2009, Avila et al. 2012, Rezával et al. 2012, Rezával et al. 2014). In addition to the five neural genes we identified from the initial test, three other genes had been reported to modulate sperm competition outcome through *ppk+* neurons (*para, Rab2, Rim*; Chow et al. 2013). Therefore, in the secondary test, we assessed the effect of knocking down each of these eight genes in *Tdc2*^+^ neurons and *ppk*^+^ neurons.

Of these eight genes, *caup* was the only gene that affected P1 when knocked down in *Tdc2*^+^ neurons. Knockdown females produced much fewer first male progeny and slightly more second male progeny than control females over the course of the assay, resulting in an overall reduction in fertility and significant decreases in P1 in vials 2-5 (Fig. 1K). *Hid*, one of the genes that affected P1 when knocked down pan-neuronally, had no overall effect on P1 when knocked down in *Tdc2*^+^ neurons. However, on a vial by vial basis, P1 in vials 2 and 3 was significantly higher in *hid* knockdown females than that of controls, under *Tdc2*^+^ specific knockdown (Fig. 1L). This result suggests a weaker, but significant role for *hid* in *Tdc2*^+^ neurons on sperm competition outcome.

We also corroborated earlier findings and showed that *ppk*^+^ neuron-specific knockdown of *Rim* caused females to produce fewer first male progeny and more second male progeny than control females while keeping total progeny constant, thus leading to lower P1 (Fig. 1M). None of the other seven genes affected P1 when knocked down in *ppk*^+^ neurons. This included *Rab2* and *para*, two genes that had been reported to affect P1 upon knockdown in *ppk+* neurons by Chow et al. (2013). That study used a different *ppk*-GAL4 driver from our study, possibly explaining the discrepancy; alternatively, variable environmental factors could be the cause.

## Discussion

A number of approaches have suggested that females play an active role in sperm competition, but the genes and cell types that mediate this have not been determined. Here, we identified such genes and also showed that their action in specific neurons is required, indicating an active role for the female in sperm competition. We functionally tested 27 candidate genes detected in a GWA study (Chow et al. 2013) for their contribution to the female’s mediation of sperm competition. We found that knockdown of eight genes in females affected P1. Six of eight mediated a change in P1 when knocked down in the female’s nervous system. Knockdown of the remaining 19 genes tested either had no detectable effect on sperm competition (perhaps another gene near the SNP is involved), or their role in sperm competition could not be identified given limitations of the RNAi method. Specifically, there could be functional redundancy, or, because the SNPs identified in Chow et al. (2013) were mostly in non-coding regions, the SNPs could affect the level of gene expression in a way that could not be probed by simple quantitative knockdown.

Earlier studies identified two genes in females (*SPR* and *Neprilysin 2*) that affected sperm competition when knocked down ubiquitously, and that are expressed in the female nervous system among other tissues (Chow et al. 2010, Sitnik et al. 2014). However, in addition to identifying new genes that, in females, regulate sperm competition outcomes, our study goes further, by directly knocking down genes in the female nervous system and showing that this affects the paternity success of competing males.

Several aspects influence which male sires most offspring in a competitive context. Sperm competition experiments using males with fluorescently labeled sperm protamines have shown that the number of sperm transferred by either male and sperm displacement by a competitor play a role (Manier et al. 2010). On the female side, her receptivity to repeated matings, influenced by her detection of male courtship and pheromones (Smith et al. 2017), directly impact the risk and intensity of sperm competition. Furthermore, the sperm ejection time after second mating affects which sperm are stored and therefore contributes to the fertilization set (Manier et al. 2010, Lüpold et al. 2013). The *Dh44*^+^ neural circuitry controls sperm ejection (Lee et al. 2015), but to our knowledge, it has not been tested whether Dh44 or *Dh44*^+^ neurons play a direct role in sperm competition. On the other hand, uterine conformational changes modulated by muscle contractions have been shown to affect sperm storage (Adams and Wolfner 2007, Mattei et al. 2015). In addition, maintenance of sperm viability once in storage (Schnakenberg et al. 2012) and detection of seminal fluid proteins like sex peptide (SP; Chapman et al. 2003, Liu and Kubli 2003) could affect sperm competition outcome as well. Finally, at least in *D. melanogaster*, there is evidence for the fair raffle hypothesis, which suggests that once sperm is in storage, there is an equal chance for each sperm to be used, regardless of which male provided the sperm (Parker 1990, Manier et al. 2010).

It is conceivable that our gene knockdowns impact neuronal signaling and consequently female physiology, behavior, or muscle contractions, allowing for any of these female-mediated aspects of sperm competition to be affected. In line with this hypothesis, we identified a role for both sensory *ppk*^+^ neurons and octopaminergic *Tdc2*^+^ neurons in mediating sperm competition outcome. A population of sexually dimorphic *Tdc2*^+^ neurons located at the posterior tip of the abdominal ganglion innervate the female reproductive tract extensively and regulate post-mating responses (PMR) including remating refractoriness and ovulation (Rezával et al. 2014). *Tdc2*^+^ neuronal innervation of the female sperm storage organs (seminal receptacle and paired spermathecae; Avila et al. 2012, Rezával et al. 2014) suggests the hypothesis that *caup*, which has a basic role in neuronal development (Gómez-Skarmeta et al. 1996), or *hid*, with its role in apoptosis (Grether et al. 1995) might affect development of *Tdc2*^+^ neurons, which in turn could modulate sperm storage and sperm competition. In addition, sensory *ppk*^+^ neurons are also crucial for female PMR (Häsemeyer et al. 2009, Yang et al. 2009). The male seminal fluid protein SP binds to the SPR expressed in female *ppk+* neurons to silence these neurons and elicit PMR (Yapici et al. 2008, Häsemeyer et al. 2009, Yang et al. 2009, Rezával et al. 2012, Lee et al. 2016). Both SP and SPR are known to influence sperm competition outcome (Chow et al. 2010, Castillo and Moyle 2014). *Rim’s* general function in the nervous system is to mediate efficient neurotransmitter secretion (Graf et al. 2012, Müller et al. 2012). Thus, *Rim* knockdown in the *ppk*^+^ neurons could affect these neurons’ signaling capabilities, thereby mediating a change in P1. SP and SPR silence the *ppk+* neurons to induce increased egg production and lower re-mating rate. In this regard, it might be surprising that *Rim* knockdown does not mediate these PMR. However, all females in our experiments are mated and thus exposed to SP, so the effect of *Rim* knockdown in a mated female might not have extra effects on PMR in addition to the *ppk*^+^ neuron-silencing effects SP already has. Finally although the female reproductive tract is, itself, extensively innervated, seminal fluid proteins can also enter the female’s hemolymph (Monsma et al. 1990, Lung and Wolfner 1999, Ravi Ram et al. 2005, Pilpel et al. 2008) and thus have the opportunity to directly interact with *Tdc2*^+^ or *ppk*^+^ neurons throughout the female body.

In addition to *caup, hid* and *Rim*, the products of three additional genes for which we found a sperm competition function might affect functioning of the female nervous system: Msp300 has previously been found to play a role at the neuromuscular junction (Morel et al. 2014); Btsz is a synaptotagmin-like protein involved in membrane trafficking (Serano and Rubin 2003); and Ddr belongs to the family of receptor tyrosine kinases, but its exact function is unknown (Sopko and Perrimon 2013).

Two of the eight genes that affected P1 were only tested with ubiquitous knockdown. *CG32834*, a predicted serine-type endopeptidase, is female spermathecae-specific (Leader et al. 2018). The spermathecae are long-term sperm storage organs whose secretions affect sperm motility (Schnakenberg et al. 2011), so *CG32834* has the potential to affect sperm storage, maintenance or release from storage. In addition, a previous study found that *CG32834* knockdown results in lower egg production and increased re-mating (Sirot et al. 2014a), in line with results reported here. *CG31872* is reported to be expressed in the female rectal pad (Leader et al. 2018), but it is not clear what its function is in female reproduction. These can be further investigated in the future by testing tissue-specific knockdowns.

Across all tissues tested, six of the eight genes that affected sperm competition outcome when knocked down also led to a decreased success for the first male. This suggests that in a wild-type situation, these genes, or the neurons in which they act, normally play a role in mediating a higher paternity success for the first male. It is possible that, when these genes are knocked down, neuronal signaling in response to the first mating is impaired. This could lead to decreased storage of first male’s sperm, increased loss or displacement of first male’s sperm, or an incomplete switch from virgin to mated state. This could also explain the lower overall fertility that we often observed in knockdown females. A second mating, and a second exposure to mating signals, mechanical and/or molecular, might improve the response to mating, leading to a higher success for the second male. Since the candidate genes tested here were identified based on natural variation in the DGRP, where females from some isofemale lines naturally have a lower P1 when doubly mated to standard tester males (Chow et al. 2013), it is possible that there is natural variation in how strongly females respond to mating due to variation in neural development, or differences in neural gene expression. All SNPs in the 33 candidate genes identified by Chow et al. (2013) are in noncoding regions or are synonymous substitutions, suggesting that they can indeed affect gene expression. Durham et al. (2014) measured variation in fecundity across young and aged DGRP females mated to males of their own line and identified associated candidate genes in a GWAS. GO categories overrepresented in those GWAS results included categories associated with neural development (Durham et al. 2014), and five of their candidate genes were also found in the GWAS from Chow et al. (2013) (*Ddr, CG32834, sima, Rbp6* and *CG15765).* Genotype-specific differences in fecundity could exist because the optimal number and timing of egg production can be a source of sexual conflict: it is beneficial for males if a female produces many eggs shortly after mating (and before remating), while more reserved resource allocation can be more beneficial for females (Sirot et al. 2014b, Wensing and Fricke 2018).

Finally, an outstanding question in research on sexual conflict is concerned with the interplay of male signals that act on the female’s nervous system to influence her physiology and behavior, and the female’s processing of and response to male cues (Schnakenberg et al. 2012). Our findings regarding sensory *ppk*^+^ neurons and *Tdc2*^+^ neurons, which include motor neurons innervating the female reproductive tract, form an important step in understanding the mechanistic and molecular basis of that interplay in sperm competition.

## Acknowledgements

We thank members of the Clark and Wolfner labs for helpful discussions and comments. We are grateful to NIH R01 HD059060 (to AGC and MFW) for supporting this work. Several authors were supported for all or part of this study by fellowships, for which we are grateful: a SUNY Diversity Fellowship, followed by an HHMI Gilliam Fellowship (SLW), a Belgian American Educational Foundation (SYND), and a summer internship from the NSF-funded EDEN Research Collaboration Network (C. Extavour, PI) for MC.

